# Opposite Effects of Alpha Oscillations on Mind-wandering With Eyes Open and Closed

**DOI:** 10.64898/2026.01.20.700111

**Authors:** Esther M. Thielking, Hannah Ramsey, Jason Samaha, Thomas Andrillon, Luca Iemi

**Author notes:** **Corresponding author**: Esther M. Thielking.

## Abstract

Spontaneous fluctuations in brain activity are thought to underlie mind-wandering, moments when mental experience becomes unrelated to what one is currently doing. A large body of research has investigated ongoing neural oscillations in the alpha band (7-14 Hz), reflecting changes in arousal and attention. However, studies disagree whether the power of alpha oscillations increases or decreases during mind-wandering compared to on-task focus. We hypothesized that these opposite effects arise from differences in eye state across studies, as eye closure increases alpha power and can reverse alpha’s relationship with sleepiness, which is itself related to mind-wandering. To test this, we recorded EEG while male and female participants with eyes either open or closed were probed to report whether they were focused on the auditory attention task or mind-wandering, as well as how sleepy they were. Consistent with our hypothesis, we found that increased alpha power made both mind-wandering and sleepiness more likely in the eyes-open group but less likely in the eyes-closed group. A systematic review of past literature largely replicated these opposite relationships across eye states in different paradigms. We propose that these results reflect an inverted-U relationship between alpha power and mind-wandering, formed by a positive relationship at low alpha power values typical of eyes-open states and a negative relationship at high alpha power values typical of eyes-closed states. By sampling alpha power across eye states in the same paradigm, this work reconciles contradictions in prior literature and clarifies how ongoing neural oscillations reflect the stream of mental experiences.

**Significance Statement:** Brain signals tracked with electroencephalography (EEG) can provide a read-out of a person’s current mental experience. Increases in a brain signal called alpha oscillations usually indicate sleepiness when recorded with eyes open. Surprisingly, the read-out can be reversed when eyes are closed: increases in alpha oscillations indicate alertness. Here, we found that this reversal explains the puzzling relationship between alpha oscillations and mind-wandering, when mental experience becomes unrelated to the task-at-hand. In moments of increased alpha, participants with eyes open reported more mind-wandering and sleepiness, whereas participants with eyes closed reported more on-task focus and alertness. One intriguing possibility is that decreases in arousal reflected by changes in alpha oscillations trigger episodes of mind-wandering.

## Introduction

Our mental experience can spontaneously become unrelated to what we are currently doing and perceiving: this is the common experience of mind-wandering. At the same time, the state of the brain changes from moment to moment due to ongoing, internally generated neural processes occurring independently of task demands or sensory input. How do these spontaneous fluctuations of mental experience and brain state relate to each other? To address this question, numerous studies have recorded a prominent type of spontaneous brain activity, alpha oscillations (7–14 Hz), while participants sustained their attention on repetitive tasks and intermittently reported mind-wandering (Kam et al., 2022). These studies show that moment-to-moment fluctuations in alpha power are correlated with mind-wandering reports. However, the direction of this relationship has been inconsistent: some studies report positive relationships (stronger alpha power while mind-wandering; Compton et al., 2019; Arnau et al., 2020; Nakatani et al., 2024), whereas others report negative relationships (weaker alpha power while mind-wandering; Braboszcz and Delorme, 2011; van Son et al., 2019a; Rodriguez-Larios and Alaerts, 2021; Rout et al., 2024).

These findings have been described as “conflicting,” “inconsistent,” or “complex” (Compton et al., 2019; Arnau et al., 2020; Kam et al., 2022; Nakatani et al., 2024), and although methodological differences have been proposed as an explanation, the debate remains unresolved. We reasoned that these inconsistencies reflect differences in eye state across studies. Closing the eyes increases the power of alpha oscillations (Berger, 1929) and can reverse the direction of its relationship with arousal (e.g., subjective sleepiness: Strijkstra et al., 2003; Kaida et al., 2006), which is thought to influence mind-wandering (Andrillon et al., 2019; Jubera-García et al., 2021). Accordingly, we propose an underlying inverted-U function relating mind-wandering with alpha power across eye states. Here, mind-wandering and alpha power are positively correlated when alpha oscillations fluctuate within a low power range as during eyes-open states, corresponding to the left side of the inverted-U (Fig. 1). By contrast, mind-wandering and alpha power are negatively correlated when alpha oscillations fluctuate within a high power range as during eyes-closed states, corresponding to the right side of the inverted-U (Fig. 1). In this way, the seemingly contradictory positive and negative relationships observed in prior literature can be understood as reflecting different sides of an underlying non-linear function.

**Figure 1.**
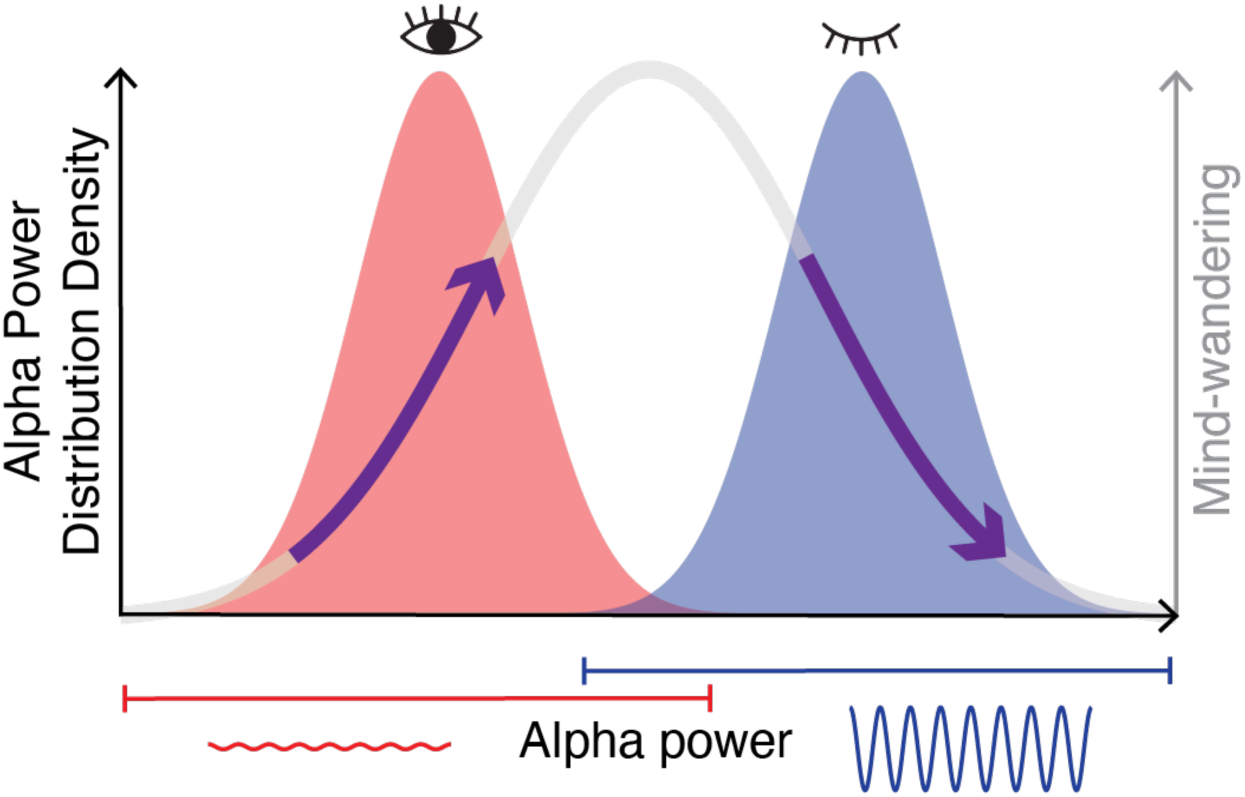
Inverted-U model. Schematic illustration of the inverted-U relationship between alpha (7-14 Hz) power and mind-wandering. The x-axis represents the range of alpha power values observed across eye states. The left y-axis shows the distribution density of alpha power values. The red distribution represents eyes-open states corresponding to the low alpha power range, while the blue distribution represents eyes-closed states corresponding to the high alpha power range. The right y-axis shows mind-wandering rate, with the gray curve depicting the inverted-U relationship between alpha power and mind-wandering rate: the left side spanning the low alpha power range shows a positive relationship, and the right side spanning the high alpha power range shows a negative relationship.

To test this, we recorded EEG during eyes-open and eyes-closed resting states followed by a sustained attention-to-response task (SART; auditory go/no-go) with intermittent experience-sampling probes and subjective sleepiness ratings. This paradigm allowed us to quantify fluctuations in mind-wandering and sleepiness through subjective reports, as well as reaction times (RTs), known to co-vary with mind-wandering (Leszczynski et al., 2017; Andrillon et al., 2021). To sample different sides of the inverted-U function, we manipulated the range of alpha power values by varying eye state between participants: one group performed the task with eyes open (EO group), while another group performed it with eyes closed (EC group). By comparing alpha power between the task and the resting states, we confirmed that eye-closed states boosted alpha power and thus produced a high power range relative to eyes-open states (Barry et al., 2007). For each group, we then assessed how ongoing alpha power was correlated with momentary fluctuations of subjective experience and RTs. We found that alpha power showed a positive correlation with subjective reports of mind-wandering and sleepiness in the EO group, and a negative correlation in the EC group, consistent with the inverted-U model.

Further, a systematic literature review dividing the studies by eye state largely replicated this pattern of positive and negative relationships, further supporting the inverted-U model. RTs did not co-vary with mind-wandering and were negatively correlated with alpha and beta power only in the EC group, replicating previous results obtained in the same eye state. Taken together, the results of this study reconcile inconsistencies in the literature and help specify the link between spontaneous neural oscillations and ongoing mental experience.

## Materials and Methods

### Participants

Sixty healthy adults participated in this study. All provided written informed consent prior to data collection. Participants were randomly assigned to perform the main task with eyes either closed (EC group; N=30) or open (EO group; N=30). Before pre-processing, participants were excluded due to EEG acquisition issues (N=5) or insufficient probe response variability (i.e., percentage mind-wandering reports below 10% or above 90%; N=5). Groups did not differ in sex (EO: 4M 22F, EC: 1M 23F; Fisher’s exact test, p=0.351) or age (EO: mean 19.8 ± standard error of the mean (SEM) 0.23 years; EC: mean 19.7 ± SEM 0.19 years; t_48_=0.574, p=0.569). All analyses included 24 EC and 26 EO subjects. Experimental procedures were approved by the Institutional Review Board (IRB) at Barnard College.

### Behavioral Paradigm

Participants completed a resting-state and a task session (Fig. 2). The resting-state session consisted of two 2-min blocks (eyes-open and eyes-closed) presented in randomized order and separated by a short self-paced break. This was used to establish each participant’s low (eyes-open) and high (eyes-closed) ranges of alpha power values. Participants were instructed to stay still, fixate on a central mark (in eyes-open rest), and avoid thinking of anything in particular. Following the resting state session, participants completed six 5-min task blocks of an auditory sustained attention to response task (SART), each separated by a self-paced break. For the duration of each task block, participants in the EO group were instructed to maintain fixation on a cross, whereas participants in the EC group were instructed to maintain eye closure.

**Figure 2.**
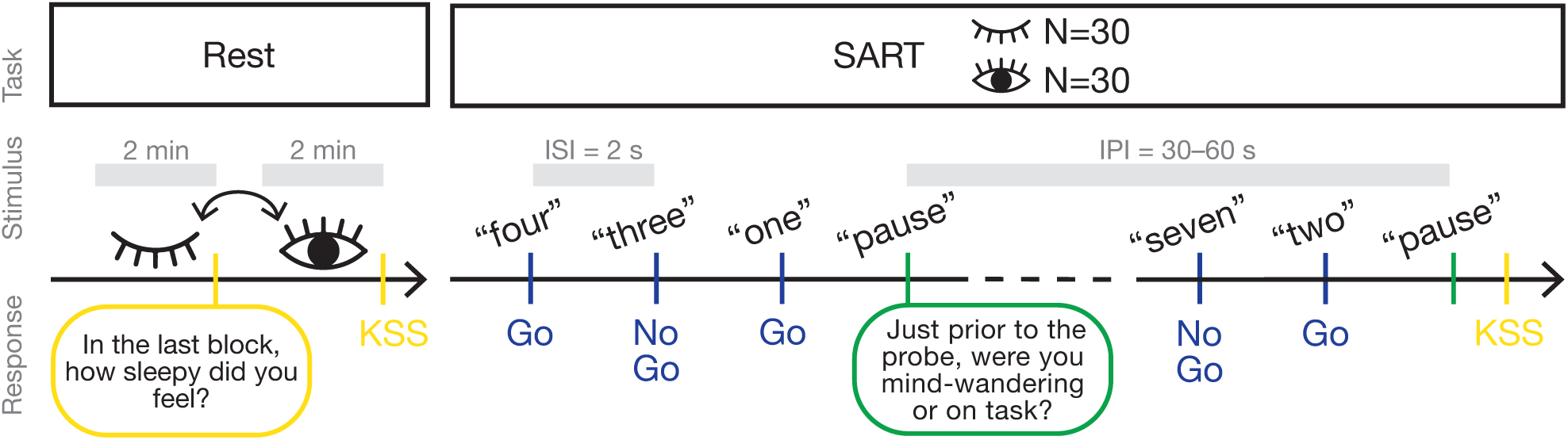
Experimental paradigm. Participants (N=60) completed an eyes-open and an eyes-closed resting state block, with each block lasting 2 min and the order randomized across participants, followed by an auditory sustained attention to response task (SART). Participants were randomly assigned to perform the SART with eyes either open or closed. The SART comprised 6 blocks, with each lasting 5 min. A spoken digit was presented on each SART trial, with an inter-stimulus interval (ISI) of 2 s: participants responded to frequent go digits and withheld responses to infrequent no-go digits. With a pseudorandom inter-probe interval (IPI) of 50 to 70 s, participants were probed to report whether they had just been on-task or mind-wandering. Sleepiness ratings (Karolinska Sleepiness Scale, KSS) were collected after each rest and task block. EEG was recorded from 8 occipito-parietal electrodes.

In the main task, participants performed an auditory SART that required them to respond as quickly and as accurately as possible via button press to frequent ‘go’ stimuli and to withhold responses to infrequent ‘no-go’ stimuli (i.e., auditory go/no-go task; Fig. 2). Task performance was measured as misses (no button press in go trials), false alarms (button press in no-go trials), and RTs (analyzed on correct go trials only). At pseudorandom intervals, participants were probed to report whether they were mind-wandering or on-task just before the probe. Participants were instructed that mind-wandering consisted of attending to task-unrelated thoughts and that on-task consisted of attending to the auditory SART (Smallwood and Schooler, 2015; Arnau et al., 2020; Compton et al., 2024). Participants were also informed that we consider mind-wandering to be a fundamental part of human cognition and were encouraged to respond honestly based on their subjective experience.

After each resting and task block, participants rated their subjective sleepiness on the 9-point Likert-type Karolinska Sleepiness Scale (KSS; 1= “extremely alert”, 9= “very sleepy, great effort to keep awake”; Kaida et al., 2006) and reported whether their eyes were open or closed to confirm compliance with the task instructions.

### Stimuli

The task was implemented in MATLAB (R2022b; The MathWorks; RRID:SCR_001622) using Psychtoolbox-3 (RRID:SCR_002881; Brainard, 1997). Auditory stimuli were synthesized with Google Cloud Text-to-Speech (*Cloud Text-to-Speech API v1*, n.d.). Spoken digits (0.499 s duration) were SART stimuli (go and no-go), and the spoken word “pause” was the experience-sampling probe (0.545 s duration). Files were sampled at 48 kHz and delivered via in-earbuds at a fixed volume. There were 9 spoken digits, ranging from “one” to “nine;” “three” and “seven” were no-go stimuli, while the others were go stimuli. Each task block included 158 trials (34 no-go, 119 go, 5 probes). The stimulus sequence was pseudorandomized for each block: 17 independent random permutations of the spoken digits were concatenated to form the base digit list (length 153) from which 5 probe insertion indices were drawn separated by a uniformly random step of 25–35 trials (mean 30.19 ± SEM 0.04 trials), corresponding to a 50-70 second inter-probe interval (IPI) comparable to previous probe-caught experience-sampling paradigms (Compton et al., 2019; Schubert et al., 2020; Andrillon et al., 2021; Rodriguez-Larios and Alaerts, 2021; Kucyi et al., 2024). The inter-stimulus-interval (ISI) was 2 s (Bastian and Sackur, 2013; Leszczynski et al., 2017).

### EEG Recording and Preprocessing

An Enobio-8 system (Neuroelectrics) was used to record continuous EEG sampled at 500 Hz. We used 8 gel channels over occipital and parietal areas of interest (P3, Pz, P4, PO3, PO4, PO7, PO8, Oz; 10-10 system), corresponding to the scalp location of the most prominent alpha rhythm in the human EEG (Tenke et al., 2015; Hohaia et al., 2022). EEG data were referenced to an ear-clip electrode. The FieldTrip toolbox (version 20180801; Oostenveld et al., 2011) running on MATLAB (R2024b; The MathWorks; RRID:SCR_001622) was used to process and analyze the data. Continuous data were epoched into 1000 ms epochs consecutively for resting-state blocks or from −2000 to 500 ms relative to stimulus onset for task blocks. Epoched data were visually inspected to remove major artifacts. One task block was removed from a participant in the EO group, who reported closing their eyes during the block. A mean of 20.94 ± SEM 5.55 epochs were rejected per participant. In three participants, one channel was excluded; in one participant, two channels were excluded.

### Spectral analysis

#### Milliseconds-scale spectral analysis

We computed time–frequency representations (TFRs) of power from the epoched data using an adaptive sliding-time window of three cycles in length (Δt = 3/f), with a Hanning taper applied before obtaining power via a fast Fourier transform (FFT) approach. The TFRs were computed at 100 ms intervals across each epoch, and spectral power was estimated from 5 to 30 Hz in 0.5 Hz steps. This milliseconds-scale TFR was used in the analysis of the relationship between spectral power and RTs.

#### Seconds-scale spectral analysis

We extracted 1000 ms consecutively for resting epochs and from −1000 to 0 ms relative to stimulus onset on each trial for task epochs, then estimated spectral power using a Hanning-tapered FFT (5–30 Hz, 0.5 Hz steps). The second half of the 2s ISI were analyzed to measure “ongoing” power, as this window minimizes contamination from evoked or induced activity related to stimulus presentation or motor responses from the preceding trial. We then constructed a seconds-scale time–frequency representation (TFR) by stacking the power spectra from the five 1-s epochs preceding each probe. This created a temporal axis of sequential pre-probe periods (e.g., −9 to −8 s, −7 to −6 s, −5 to −4 s, −3 to −2 s, and −1 to 0 s relative to probe onset) consistent with the time range analyzed in prior studies (Baldwin et al., 2017; Compton et al., 2019; Jin et al., 2019; Schubert et al., 2020; Andrillon et al., 2021; Kucyi et al., 2024). This seconds-scale TFR was used for the analysis of the relationship between spectral power and mind-wandering. We averaged the seconds-scale data within each block for the analysis of the relationship between spectral power and sleepiness.

Epoch-wise power values were averaged across channels for all analyses. Because we were primarily interested in brain activity before experimental events (pre-probe and prestimulus), we did not apply baseline correction in either the milliseconds-scale or seconds-scale spectral analyses.

### Testing the Inverted-U Model

By “inverted-U” we specify that the relationship between alpha power and subjective reports changes in direction (sign) along the range of alpha power values observed across eye states from positive sign in the low range of alpha power to negative sign in the high range of alpha power. The eye state manipulation in our study allowed us to sample low (EO group) and high (EC group) ranges of alpha power values, corresponding to the left and right sides of the hypothesized inverted-U. To test this, we fit separate linear relationships within the EO and EC groups and tested whether they have individually significant slopes with opposite signs (“two-lines” test: Simonsohn, 2018). We chose this method over fitting a single quadratic function because it is compatible with our between-subject design and because quadratic models can both make unjustified assumptions about the function’s shape and inflate false-positive rates (Simonsohn, 2018). Evidence for an inverted-U is obtained if two criteria are met: (i) eyes-open and eyes-closed states exhibit non-overlapping alpha power ranges (see *Analysis of Spectral Power and Eye State*); (ii) the EO group shows a significantly positive relationship and the EC group shows a significantly negative relationship, between alpha power and subjective reports (see *Analysis of Behavioral Responses and Their Relationship With Spectral Power*).

### Analysis of Spectral Power and Eye State

To demonstrate that eyes-open and eyes-closed states produce low and high ranges of alpha power values, respectively, we performed two analyses. First, we computed the difference in spectral power between (i) eyes-open and eyes-closed epochs from resting blocks across all participants and (ii) epochs from resting blocks (either eyes-open or eyes-closed) and an equal number of randomly selected epochs from task blocks of the opposite eye state, separately for each participant group. Second, we examined whether eyes-open and eyes-closed states were distinguished by partially non-overlapping ranges of power distributions. For each participant and frequency we computed the percentage of epochs whose power values are observed in one eye state but never in the other. The eyes-open non-overlap was defined as the proportion of eyes-open epochs with power values below the minimum power value observed in the eyes-closed distribution. The eyes-closed non-overlap was defined as the proportion of eyes-closed epochs with power values above the maximum power value observed in the eyes-open distribution. We computed these within-participant non-overlap metrics in two ways: (i) by comparing eyes-open and eyes-closed epochs from resting blocks across participants, and (ii) by comparing all epochs from resting blocks (either eyes-open or eyes-closed) and an equal number of randomly selected epochs in task blocks of the opposite eye state, separately for each participant group. At the group level, we assessed the significance of eyes-open and eyes-closed non-overlap metrics using separate cluster-based permutation tests across frequencies (see *Statistical Analyses*).

### Analysis of Behavioral Responses and Their Relationship With Spectral Power

For each participant, we used separate generalized linear models (GLMs) to examine first (i) the contribution of sleepiness ratings in explaining block-average mind-wandering probability, and second the contributions of EEG spectral power during the task in explaining (ii) mind-wandering reports, (iii) sleepiness ratings, and (iv) RTs.

For GLMs (i), (iii), and (iv), we fit standard linear regression models of the form:

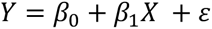

where *X* is the z-scored predictor (sleepiness rating or spectral power, the latter normalized by log₁₀-transformation: Smulders et al., 2018), *Y* is the corresponding z-scored behavioral measure (mind-wandering probability, sleepiness rating, or RTs, the latter log₁₀-transformed), *β*_0_is the intercept, *β*_1_ is the slope coefficient, and *ɛ* is the residual error. From these models, we use *β*_1_ to quantify the contribution of *X* to explaining variation in *Y*.

For GLM (ii), we fit a logistic regression model of the form:

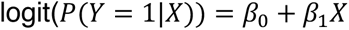

where *X* is EEG spectral power (log₁₀-transformed and z-scored), *Y* is the binary report to the experience-sampling probe (*Y*=1 for a mind-wandering report, *Y*=0 for an on-task report), and *P*(*Y* = 1|*X*) the probability of reporting mind-wandering given *X*. From this model, we use *β*_1_to quantify the contribution of EEG spectral power to explaining the log-odds of reporting mind-wandering.

### Statistical Analyses

A two-stage statistical approach was applied to test the significance of the observed statistics described in the preceding sections. At the subject level, we converted the observed statistics to z-scores using a permutation-based null distribution. This process involved randomly shuffling either the eye state (differences and non-overlap scores) or the mapping between the predictor and the behavioral measure (GLM *β*_1_) for 1000 iterations and recomputing the statistic for each permutation to create a null distribution. The observed statistic was then standardized by subtracting the mean and dividing by the standard deviation of this null distribution, resulting in one z-score for each subject, and for each frequency and time point (if applicable) (Samaha et al., 2017; Iemi and Busch, 2018).

At the group level, we tested whether the subject-level z-scores were significant across participants (i.e., whether their signs were consistent). For analyses without multiple comparisons (behavioral GLM), significance was assessed using a two-tailed t-test against zero on the subject-level z-scores. For other analyses (EEG GLMs), significance was assessed using a nonparametric cluster-based permutation test, which addresses multiple comparisons across frequencies and time points (if applicable) (Maris and Oostenveld, 2007). We obtained a distribution of z-scores under the null hypothesis by randomly permuting their signs 1000 times. On each iteration, we tested the resulting z-scores with a two-tailed t-test against zero and assessed the sum of the t values within the largest contiguous cluster of significant time-frequency points (or frequency points) (cluster alpha: p<0.05), resulting in a distribution of t sums expected under the null hypothesis. A final p-value was calculated as the proportion of t sums under the null hypothesis larger than the sum of t values within clusters in the observed data. Thus, p-values smaller than 0.05 (final alpha) indicate that the observed coefficients were significantly different from zero.

We quantified evidence for the null hypothesis using Jeffreys–Zellner–Siow (JZS) Bayes factors (BF_10_) (MATLAB BayesFactor toolbox; Krekelberg, 2021) with the default Cauchy prior on standardized effect sizes with scale r = 0.707 (Rouder et al., 2009). We interpreted BF as follows: BF > 3 indicates evidence favoring the alternative, BF < 1/3 indicates evidence favoring the null, and values between 1/3 and 3 were treated as inconclusive. For analyses with multiple comparisons that yielded a significant cluster in only one group, we extracted the mask of significant time-frequency points from that group and computed the BF at each corresponding point in the non-significant group. We then reported the percentage of time-frequency points showing evidence for the null, or inconclusive evidence (Iemi et al., 2017).

### Systematic literature review

We conducted a systematic literature review using the search engine PubMed. Our aim was to find previous studies that examined the relationship between ongoing alpha power and subjective reports of attentional state (mind-wandering) in neurotypical humans. We searched in the abstracts and titles for the following key words: (spontaneous OR pre-stimulus OR prestimulus OR ongoing OR preprobe OR pre-probe) AND (Alpha OR α OR 10 Hz OR 8–12 Hz OR 7-14 Hz) AND (variability OR oscillations OR fluctuations OR activity OR rhythmic OR power OR amplitude) AND (subjective attention OR mind-wandering OR mind wandering OR task-focused OR lapses OR misses). Inclusion required that studies analyzed neurotypical participants with no specialized training (e.g., meditators were excluded), ongoing alpha power in relation to mind-wandering reports (i.e., pre-probe window for probe-caught studies), and immediate (not retrospective) reports of attentional state. Further, we excluded studies that quantified alpha activity using a p-episode approach (e.g., Boudewyn and Carter, 2018) or that applied baseline correction to alpha power values (e.g., Hua et al., 2022).

From 29 eligible studies, we extracted (1) eye state during experience sampling: eyes open or eyes closed, (2) direction of attention: exteroceptive (attending to external stimuli) or interoceptive (attending to bodily stimuli, e.g., breath counting), with sensory modality coded for exteroceptive tasks as visual (V), auditory (A), audiovisual (AV), or (A,V) for studies with separate tasks in auditory and visual modalities, and interoceptive tasks coded as breath counting (Breath), (3) sampling method: probe-caught (experimentally-initiated report about current attentional state) or self-caught (participant-initiated reports upon recognizing mind-wandering), (4) relationship between alpha power and mind-wandering: expressed as the correlation that is expected based on the results reported by each study. In probe-caught studies, this was measured as a contrast between ongoing alpha power preceding probes where participants reported mind-wandering and on-task states; in self-caught studies, this was measured as a contrast between ongoing alpha power in the period before the button press (interpreted as mind-wandering) and after the button press (interpreted as on-task). Findings of significantly stronger alpha power before mind-wandering compared to on-task states are expressed as a positive correlation; findings of significantly weaker ongoing alpha power before mind-wandering compared to on-task states are expressed as a negative correlation; findings of no significant difference in ongoing alpha power between mind-wandering and on-task states are expressed as a “null” correlation.

### Code Accessibility

The code used for this study will be publicly released when the study is published.

## Results

### Behavior

We probed participants about once a minute to self-report whether they were focused on performing the auditory SART or engaged in task-unrelated mental experiences (i.e., mind-wandering), and about once every five minutes to report their sleepiness levels (Fig. 2). Eye state was manipulated between groups: the EO group performed the task with eyes open, and the EC group with eyes closed (Fig. 2). Participants reported frequent mind-wandering (EC 67.28±3.67%; EO 52.14±3.60%; mean ± SEM) and “some signs of sleepiness” during task performance (EO 6.083 ± 0.266; EC 6.182 ± 0.340). Mind-wandering was more likely in blocks when participants reported feeling sleepier (EO t₂₅=2.236, p=0.035; EC t_23_=4.104, p<0.001), suggesting that mind-wandering increased with sleepiness. Additionally, mind-wandering reports were preceded by more false alarms (EO t_25_=2.845, p=0.009; EC t_23_=3.845, p<0.001) and more misses over the 5 trials pre-probe in the EC group (EO t_25_=1.449, p=0.160, BF=0.524; EC t_23_=2.114, p=0.046), suggesting that mind-wandering had a detrimental effect on task accuracy. By contrast, mind-wandering didn’t affect average reaction times (RTs), demonstrated by no difference in mean RT between correct trials preceding on-task and mind-wandering reports (5 pre-probe trials: both groups p>0.517, BF<0.279; 1 pre-probe trial: both groups p>0.190, BF<0.52).

Comparing EC and EO groups allowed us to investigate the effects of eye state on participants’ behavior (Table 1). The EC group reported more instances of mind-wandering compared to the EO group, suggesting that eye closure increased mind-wandering. The effects on task performance were either null (RTs, misses) or inconclusive (false alarms, mind-wandering and sleepiness correlation). There was a greater increase in sleepiness during the task relative to rest in the EC group than in the EO group. Interestingly, across participants, eyes-closed rest was associated with higher sleepiness ratings (5.520 ± 0.220) than eyes-open rest (4.900 ± 0.259; t_49_=3.313, p=0.002), suggesting that eye closure increased sleepiness not only during the task but also during rest.

**Table 1.** Behavioral effects of eye state. Relative to the eyes-open (EO) group, the eyes-closed (EC) group showed significantly more mind-wandering and a significant increase in sleepiness during the task relative to rest. No significant between-group differences were found in the correlation (z-scored GLM β coefficient; z(β)) between mind-wandering and sleepiness (MW-S), or in reaction times (RTs), misses, or false alarms. Comparisons were made with unpaired t-tests, and Bayes factor (BF) analysis was used to determine whether non-significant results reflected evidence for the null (BF<1/3) or inconclusive evidence (1/3<BF<3).

| Behavioral Metrics | EC (mean ± SEM) | EO (mean±SEM) | $t_{48}$ | p | BF |
| --- | --- | --- | --- | --- | --- |
| Mind-wandering (%) | 67.28 ± 3.67 | 52.14 ± 3.60 | 2.944 | 0.005 | 8.384 |
| $\Delta$ Sleepiness (%) | 42.81 ± 11.47 | 11.39 ± 5.40 | 2.541 | 0.014 | 3.653 |
| MW-S correlation $z(\beta)$ | 0.795 ± 0.194 | 0.369 ± 0.165 | 1.685 | 0.099 | 0.893 |
| RTs (s) | 0.675 ± 0.021 | 0.690 ± 0.027 | -0.415 | 0.680 | 0.304 |
| Misses (%) | 1.44 ± 0.57 | 1.38 ± 0.38 | 0.090 | 0.929 | 0.284 |
| False alarms (%) | 19.80 ± 2.60 | 27.38 ± 3.56 | -1.697 | 0.096 | 0.907 |

Taken together, these behavioral results suggest that, in this study, mind-wandering is associated with increased sleepiness and worse task accuracy (but no change in average RT), and that eye closure results in increased mind-wandering and sleepiness.

### Eyes-open and eyes-closed states produce partially non-overlapping ranges of alpha power values

In this study, we used a manipulation of eye state to sample left and right sides of the proposed inverted-U function relating alpha oscillations and mind-wandering (Fig. 1). Based on prior research (Berger, 1929), we expect eyes-open states to sample the low range of alpha power values or the left side of the inverted-U, and eyes-closed states to sample the high range of alpha power values or the right side of the inverted-U (Fig. 1). Thus, we assessed criterion (i) of the inverted-U model: whether eye closure increased mean alpha power and shifted its range to include values that are not observed during eyes-open states (i.e., a different side of the inverted-U).

We first compared ongoing oscillatory power between eyes-open rest and eyes-closed rest. Cluster permutation testing combining participants from both groups revealed that eye closure during rest increased power in the alpha and surrounding bands (5–23.5 Hz, p=0.001, peak t_49_=10.869 at 10 Hz; data not visualized). We then compared ongoing oscillatory power during task engagement with that observed during rest in the other eye state (for EO and EC groups separately). Cluster-based permutation testing revealed a significant increase in power in the alpha and surrounding bands during task blocks in the EC group relative to eyes-open rest (5–24 Hz, p=0.001, peak t_23_=7.940 at 11.5 Hz; Fig. 3b), and a significant decrease in power in a similar frequency range during task blocks in the EO group relative to eyes-closed rest (6-22.5 Hz, p=0.006, peak t_25_=-6.602 at 10 Hz; Fig. 3a). Taken together, these findings confirm the well-documented “alpha blocking” effect (Berger, 1929; Barry et al., 2007; Tenke et al., 2015; ElShafei et al., 2022; Hohaia et al., 2022; Petro et al., 2022) and show that alpha power increases when the eyes are closed regardless of whether oscillatory activity is recorded during rest or task-engaged states.

**Figure 3.**
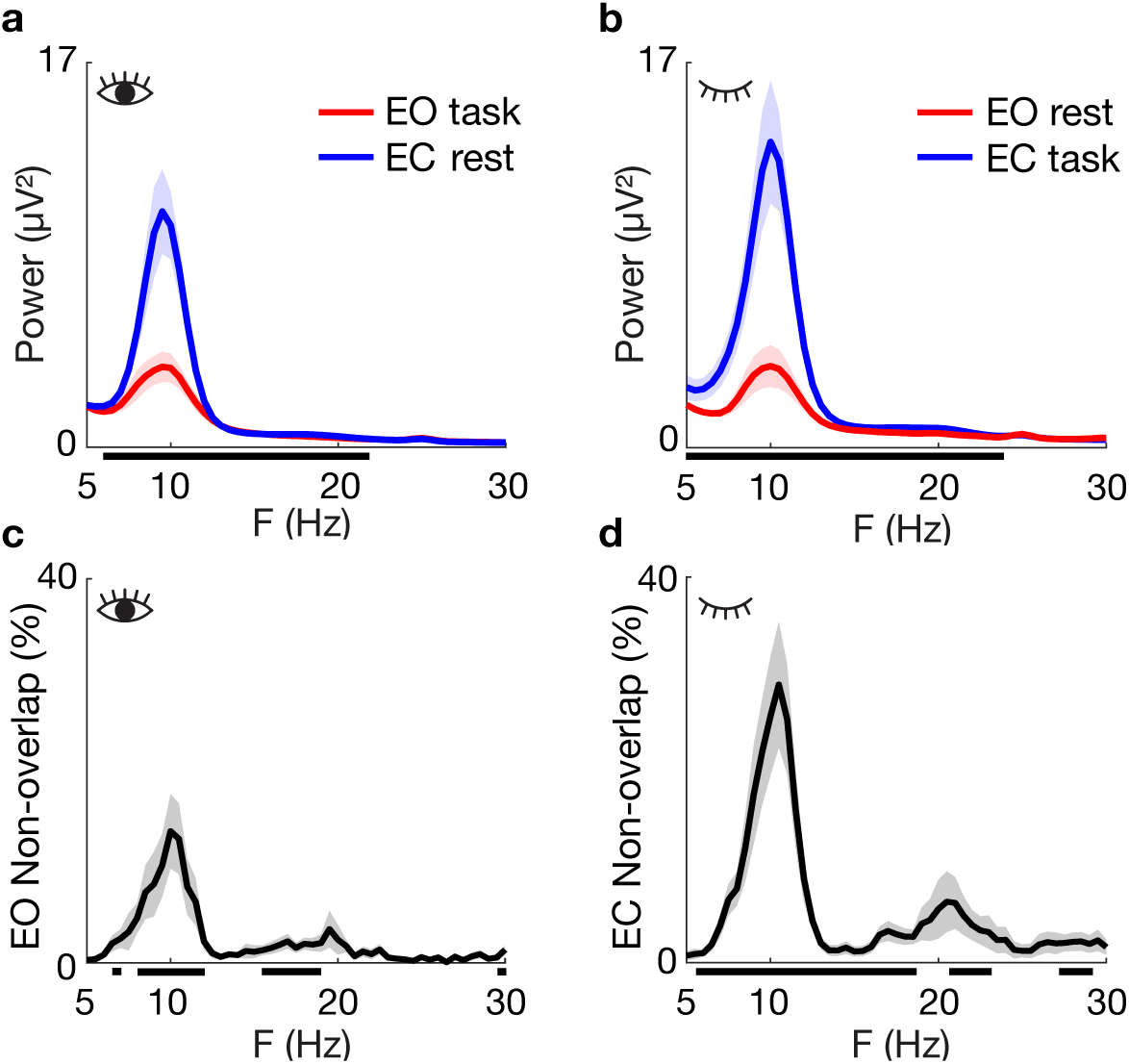
Eye-state effects on spectral power. **(a,b)** Group-average power spectra (mean ± shaded SEM) during task and eyes-closed rest in the eyes-open (EO) group (a), and during task and eyes-open rest in the eyes-closed (EC) group (b). **(c,d)** Group-average frequency representation of EO non-overlap: proportion of EO task epochs with power values below the minimum value observed in eyes–closed rest (c), and EC non-overlap: proportion of EC task epochs with power values above the maximum value observed in eyes-open rest (d). Significant frequency clusters are indicated with black horizontal lines (cluster-based permutation test, p<0.05). Each eye state is characterized by distinct, partially non-overlapping alpha power ranges: specifically, eyes-closed states increase alpha power, shifting it to values never observed during eyes-open states.

We hypothesized that manipulating eye state enables sampling alpha power in left and right sides of the inverted-U. To assess this, we measured the extent of non-overlap between the eyes-open and eyes-closed epoch-wise power distributions: the proportion of eyes-open values lower than the eyes-closed minimum (EO non-overlap) and the proportion of eyes-closed values higher than the eyes-open maximum (EC non-overlap). In resting-state data combining participants from both groups, cluster permutation testing revealed significant eyes-open non-overlap spanning the alpha band and surrounding frequencies (5.00–24.50 Hz, p<0.001, peak t_49_=4.973 at 10.5 Hz; 25.50–27.00 Hz, p=0.014, peak t_49_=3.608 at 26 Hz; data not visualized) as well as significant eyes-closed non-overlap spanning the alpha band and surrounding frequencies (5.00–13.50 Hz, p<0.001, peak t_49_=6.066 at 10.5 Hz and 16.00–22.00 Hz, p<0.001, peak t_49_=3.841 at 18.0 Hz; data not visualized). We next tested whether alpha power fluctuations during the task showed non-overlapping values with those observed during rest in the other eye state for each group separately. Cluster permutation testing revealed significant eyes-open non-overlap in the alpha and beta bands in the EO group (6.5-7 Hz, p=0.034; 8.0–12.0 Hz, p=0.001, peak t_25_=3.273 at 10 Hz; 15.5–19.0 Hz, p=0.002; 29.5-30 Hz, p=0.033; Fig. 3c) and significant eyes-closed non-overlap in the alpha and surrounding bands in the EC group (6–19 Hz, p=0.001, peak t_23_=4.96 at 12.5 Hz; 21–22.5 Hz, p=0.044, and 27.5–29.5 Hz, p=0.037; Fig. 3d).

Taken together, these results confirm criterion (i) of the inverted-U model: the eye-state manipulation produced non-overlapping alpha-power ranges corresponding to the left (eyes open) and right (eyes closed) sides of the hypothesized inverted-U function. Critically, establishing this non-overlap allows us to evaluate the relationship between alpha power and mind-wandering on each side of the inverted-U.

### Alpha power shows an inverted-U relationship with mind-wandering and sleepiness

We tested a model proposing that mind-wandering follows an inverted-U shaped function across the range of alpha power values observed in eyes-open and eyes-closed states (Fig. 1). The left side of the inverted-U corresponds to the lower range of alpha power values (sampled here during eyes-open states) where a positive correlation with mind-wandering probability is predicted; the right side of the inverted-U corresponds to the higher range of alpha power values (sampled here during eyes-closed states) where a negative correlation with mind-wandering probability is predicted. Thus, we assessed criterion (ii) of the inverted-U model by fitting separate GLMs for the EO and EC groups to examine the pre-probe, trial-wise correlation between oscillatory power and mind-wandering reports.

Cluster-based permutation testing revealed in the EO group, a significant positive correlation between mind-wandering probability and alpha power over the 5 seconds preceding the probe (11.5–13.5 Hz, –5 to 0 s, p<0.001, peak t_25_=3.904 at 12.5 Hz and 0 s; Fig. 4 a,c), and in the EC group, a significant negative correlation between mind-wandering probability and alpha power over the 7 seconds preceding the probe (9–12.5 Hz, –7 to 0 s, p<0.001, peak t_23_=–3.825 at 9.5 Hz and –7 s; Fig. 4 b,c), consistent with the inverted-U model.

**Figure 4.**
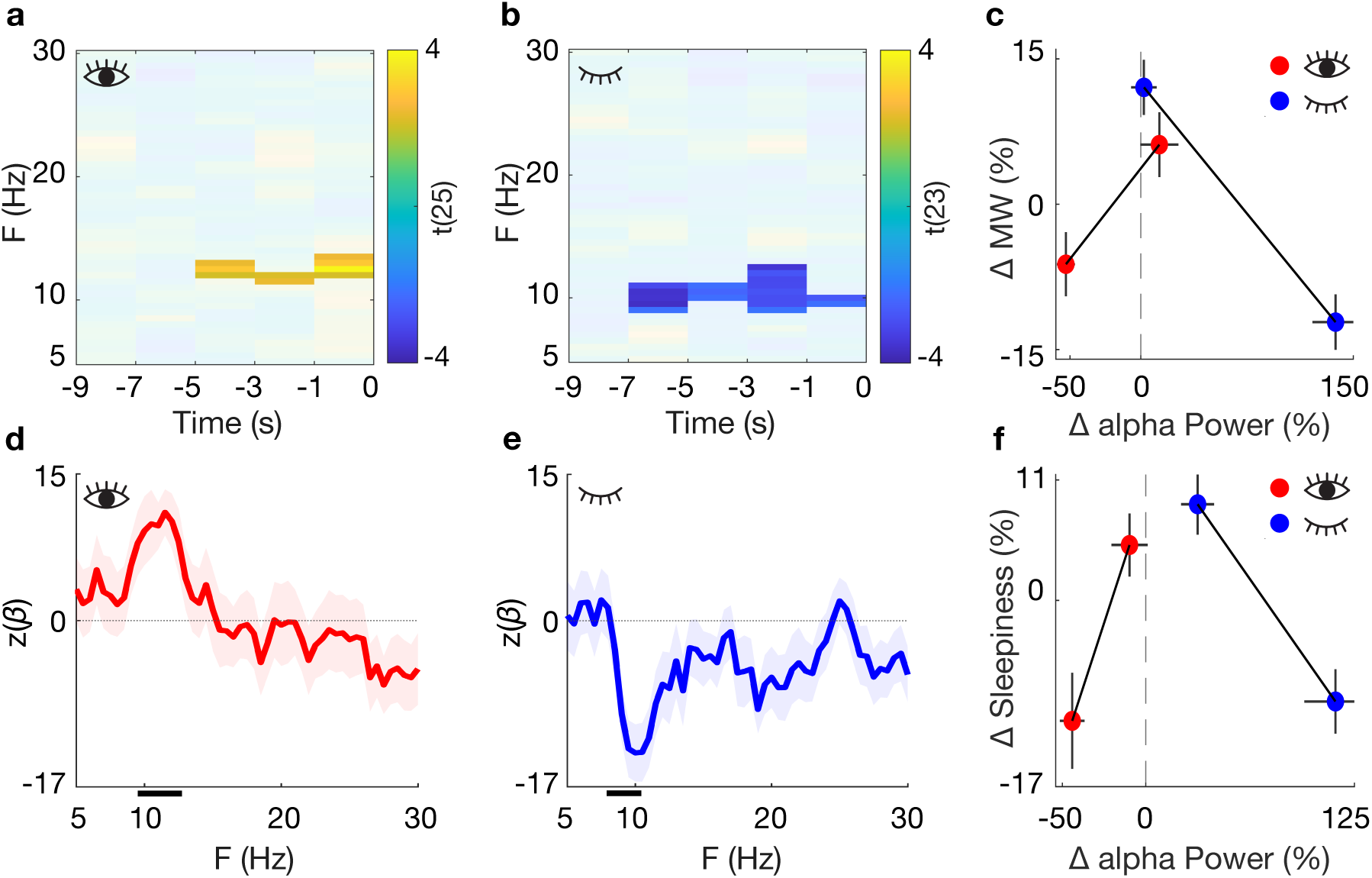
Relationship between power and subjective reports of mind-wandering and sleepiness. Participants were probed to report whether they were on-task or mind-wandering during each task block and to rate their sleepiness after each block. We used generalized linear models (GLMs) to correlate fluctuations in ongoing spectral power with each of these subjective reports separately in the eyes-open (EO) and eyes-closed (EC) groups. **(a, b)** Group-level t-statistics map the correlation between pre-probe power and the probability of reporting mind-wandering for the EO (a) and EC (b) groups, respectively. Blue indicates that mind-wandering became more likely with decreasing pre-probe power; yellow indicates that mind-wandering became more likely with increasing pre-probe power. Maps are masked to show cluster-corrected significant effects (p<0.05). Time 0 s denotes probe onset. **(d, e)** Group-level frequency-domain representations of the correlation between power and sleepiness ratings for the EO (d) and EC (e) groups. Significant frequency clusters are indicated with black horizontal lines (cluster-based permutation test, p<0.05). **(c, f)** Descriptive visualization of the inverted-U across EO and EC groups. For each participant, power was first averaged across time-frequency (c) or frequency (f) points identified as significant in either group by the cluster-based permutation tests shown in (a)/(b) for mind-wandering ((c): 9–13.5 Hz, −7 to 0 s), and in (d)/(e) for sleepiness ((f): 9–12 Hz). Trials (mind-wandering) or blocks (sleepiness) were median split into low and high power bins, and mind-wandering probability (c) or mean KSS rating (f) was computed for each bin. To represent EO and EC groups on a common axis, power and behavioral measures were expressed as percent change relative to each participant’s resting baseline (mean power across EO and EC rest) and overall mean report, respectively. Error bars show group SEM (x: power; y: behavior). Panels (c) and (f) are descriptive summaries of the results in panels (a)/(b), and (d)/(e), respectively, and were not independently tested.

Note that alpha power in the pre-probe window may be influenced by events in the preceding trial. For example, a motor response in trial t may induce alpha-band desynchronization in trial t + 1 (Pfurtscheller and Lopes Da Silva, 1999; Nikulin et al., 2007; Mazaheri and Jensen, 2008). Because mind-wandering reports are preceded by more button presses due to increased false alarm rates, changes in pre-probe alpha power attributed to mind-wandering might instead reflect differences in motor-related desynchronization, rather than endogenous fluctuations. To rule out motor confounds, we repeated the GLM analysis including only pre-probe epochs following correct go responses, all of which involved a button press in the preceding trial. This control analysis replicated the positive correlation in the EO group (p=0.021) and the negative correlation in the EC group (p<0.001), ruling out motor confounds.

The inverted-U model of mind-wandering was motivated by prior findings suggesting an inverted-U relationship between alpha power and sleepiness, a state closely related to mind-wandering (Jubera-García et al., 2021) characterized by positive and negative correlations during eyes-open and eyes-closed states, respectively (Kaida et al., 2006). To replicate these findings, we applied the same approach used for the mind-wandering analysis, fitting separate GLMs in the EO and EC groups to test the block-wise relationships between ongoing oscillatory power and sleepiness ratings. Cluster-based permutation testing revealed a significant positive correlation between sleepiness and alpha power in the EO group (9.5–12 Hz, p=0.013, peak t_25_=4.298 at 11.5 Hz; Fig. 4 d,f) and a significant negative correlation in the EC group (9–11.5 Hz, p=0.011, peak t_23_=-4.633, at 10 Hz; Fig. 4 e,f), consistent with previous results. The restricted GLMs controlling for motor confounds replicated the positive correlation in EO group (p=0.021) and negative correlation in EC group (p=0.006).

Taken together, these results confirm criterion (ii) of the inverted-U model: alpha power shows a positive correlation with mind-wandering and sleepiness at low power ranges (EO group) and a negative correlation at high power ranges (EC group), providing evidence for an inverted-U relationship between alpha power and subjective reports.

### Alpha and beta power show a negative relationship with reaction times in the eyes-closed group

Then, we examined the relationship between prestimulus alpha power and RTs to replicate previous findings (e.g., ElShafei et al., 2018). We applied the same approach as for the analysis of subjective reports, fitting separate GLMs for the EO and EC groups. Cluster-based permutation testing in the EC group showed a significant negative correlation between RTs and power in the alpha and beta bands within approximately one second before stimulus onset (6.5– 25 Hz, −1.1 to −0.3 s, p=0.018, peak t_23_=−3.593 at 19 Hz and −0.9 s; Fig. 5 b,c), indicating that stronger oscillations in alpha and beta bands over occipital and parietal scalp regions preceded faster responses to auditory stimuli. By contrast, cluster-based permutation testing in the EO group revealed no significant clusters (Fig. 5 a,c). Bayesian analysis at each time-frequency point within the EC significant cluster showed that, in the EO group, 33% of points favored the null hypothesis and 64% were inconclusive.

**Figure 5.**
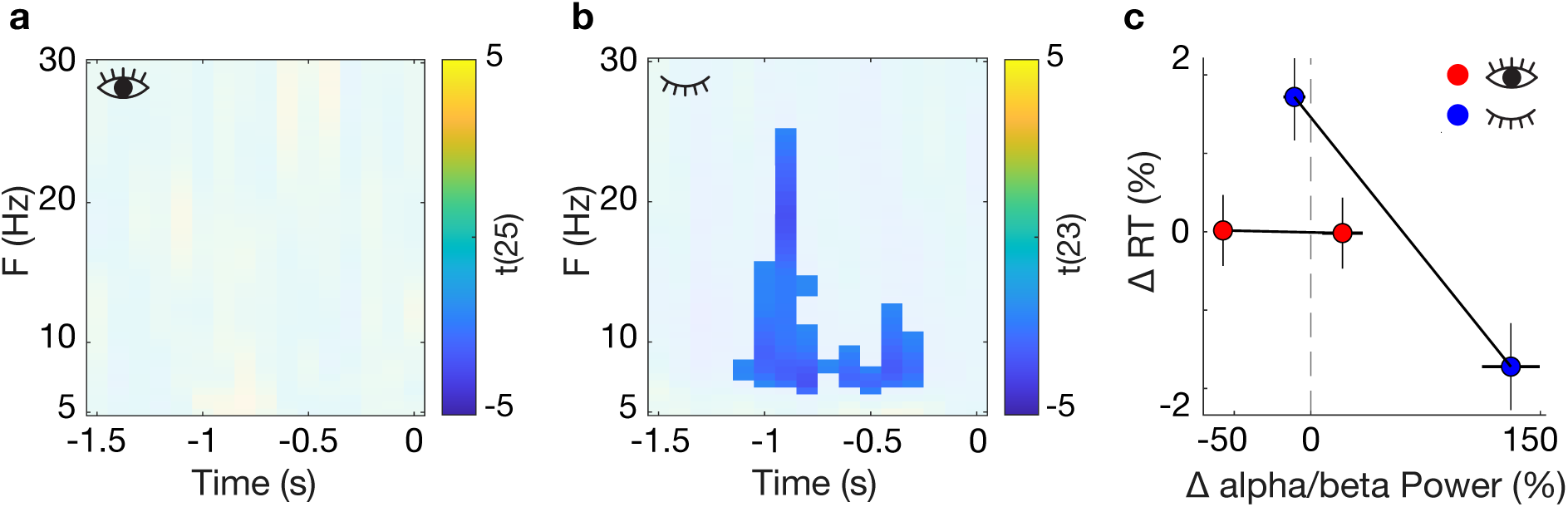
Relationship between prestimulus power and reaction times. Group-level t statistics map the correlation coefficient reflecting the relationship between EEG power and reaction times (RTs) in the (a) eyes-open (EO) and (b) eyes-closed (EC) groups. Blue and yellow values indicate a negative and positive correlation between ongoing power and RT, respectively. The maps are masked to show cluster-corrected significant effects (p<0.05). Time 0 s denotes stimulus onset. (c) Descriptive visualization of the EO and EC group comparison across resting-state normalized data. For each participant, power was first averaged across the significant points from the cluster-based permutation test of the EC group ((b): 6.5–25 Hz, −1.1 to −0.3 s). Trials were then median split into low vs high power bins, and average RT was computed for each bin. To place EO and EC groups on a common axis, power and RT measures were expressed as percent change relative to each participant’s resting baseline (mean power across EO and EC rest) and overall mean RT, respectively. Error bars show group SEM (x: power; y: RT). (c) is a descriptive summary of the results in panels A and B and was not independently tested.

To rule out motor confounds, we repeated the GLM analysis using two subsets of epochs: (i) epochs following correct go responses, which were always preceded by a button press, and (ii) epochs following correct no-go responses, which were not preceded by a button press. These control analyses replicated the inconclusive correlation in the EO group (no significant correlation in the alpha band) and the negative correlation spanning alpha and beta bands in the EC group (i: p=0.028; ii: p=0.007).

Note that average RT was not affected by mind-wandering (see *Behavioral Results*), therefore these results cannot provide evidence for or against the inverted-U model of mind-wandering. Instead, the results in the EC group replicate some previous findings showing that RTs are negatively correlated with prestimulus alpha power from task-unrelated areas.

### Systematic Literature Review Supports an Inverted-U Relationship Between Alpha Power and Mind-Wandering

We conducted a systematic literature review of studies analyzing the relationship between the power of alpha oscillations and immediate mind-wandering reports in neurotypical individuals. We categorized studies based on whether the task was performed with eyes open (Table 2, top) or eyes closed (Table 2, bottom). We found that 17/24 eyes-open studies reported a positive relationship, 3/24 reported a negative relationship, and 4/24 reported no significant relationship. This shows that prior eyes-open studies are predominantly consistent with our results in the EO group and with the left side of the inverted-U. Our study replicated this positive relationship using an auditory task, whereas most eyes-open studies (16/24) used visual tasks. Notably, although the only eyes-open study that used a self-caught methodology reported a negative relationship, alpha power in that study was higher in a distant pre-report window labeled “mind-wandering” (−10 to −5 s) than in the immediately pre-report window labeled “awareness of mind-wandering” (−5 to 0 s) (Shinagawa et al., 2025). If meta-awareness reflects an on-task state, this would constitute a positive relationship consistent with most eyes-open studies, and with the left side of the inverted-U. All 5/5 eyes-closed studies reported a negative relationship, consistent with our results in the EC group and with the right side of the inverted-U. We additionally found that 23/24 prior eyes-open studies used exteroceptive tasks and probe-caught mind-wandering reports, whereas all eyes-closed studies used interoceptive tasks and most used self-caught mind-wandering reports. Accordingly, prior literature alone could not have determined whether the opposite relationships in eyes-open and eyes-closed studies reflect different sides of the inverted-U (compatible with our model), or different mechanisms related to the type of sustained attention task or reporting method used.

**Table 2.** Review of prior studies examining the relationship between alpha power and mind-wandering. Studies using eyes-open and eyes-closed paradigms are categorized by sustained-attention task type, mind-wandering report method, and the direction of the observed relationship: “positive” means alpha power increases in mind-wandering relative to on-task states, “negative” means it decreases, and “null” means no reliable relationship. Exteroceptive tasks used visual (V), auditory (A), audiovisual (AV), or auditory and visual (A,V) modalities separately, while all interoceptive tasks involved breath counting (Breath). Probe-caught studies contrast activity preceding probes classified as mind-wandering versus on-task. Self-caught studies contrasted alpha power immediately before (mind-wandering) and after (on-task) the report. Note that one study (*) excluded the 5-s pre-probe interval to avoid conflating mind-wandering with processes related to meta-awareness. The opposite relationships observed in prior eyes-open and eyes-closed studies may reflect different sides of the inverted-U or methodological differences.

| Reference | Relationship<br>between alpha<br>power and<br>mind-wandering | Sustained attention<br>task (modality) | Mind-wandering<br>report<br>method |
| --- | --- | --- | --- |
| **Eyes-open paradigm** |  |  |  |
| Arnau et al. (2020) | Positive | Exteroceptive (V) | Probe |
| Baird et al. (2014) | Negative | Exteroceptive (V) | Probe |
| Baldwin et al. (2017) | Positive | Exteroceptive (V) | Probe |
| Broadway et al. (2015) | Null | Exteroceptive (V) | Probe |
| Compton et al. (2019) | Positive | Exteroceptive (V) | Probe |
| Compton et al. (2024) | Positive | Exteroceptive (V) | Probe |
| Conrad and Newman (2021) | Positive | Exteroceptive (AV) | Probe |
| Dhindsa et al. (2019) | Null | Exteroceptive (AV) | Probe |
| Dias Da Silva et al. (2022) | Positive | Exteroceptive (V) | Probe |
| Esposito et al. (2022) | Positive | Exteroceptive (V) | Probe |
| Gouraud et al. (2021) | Positive | Exteroceptive (AV) | Probe |
| Groot et al. (2021) | Positive | Exteroceptive (V) | Probe |
| Ibáñez-Molina and Iglesias-Parro (2014) | Null | Exteroceptive (AV) | Probe |
| Jin et al. (2019) | Positive | Exteroceptive (V) | Probe |
| Kam et al. (2021) | Positive | Exteroceptive (V) | Probe |
| Kam et al. (2025) | Positive | Exteroceptive (AV, A,V) | Probe |
| Kirschner et al. (2012) | Positive | Exteroceptive (V) | Probe |
| Macdonald et al. (2011) | Positive | Exteroceptive (V) | Probe |
| Polychroni et al. (2022) | Positive | Exteroceptive (A) | Probe |
| Prieto et al. (2021) | Negative | Exteroceptive (AV) | Probe |
| Qin et al. (2011) | Null | Exteroceptive (V) | Probe |
| Shinagawa et al. (2025) | Negative | Exteroceptive (A) | Self (*) |
|  | Negative | Interoceptive (Breath) | Self |
| Wamsley and Summer (2020) | Positive | Exteroceptive (V) | Probe |
| Yang et al. (2022) | Positive | Exteroceptive (V) | Probe |
| **Eyes-closed paradigm** |  |  |  |
| Braboszcz and Delorme (2011) | Negative | Interoceptive (Breath) | Self |
| Rodriguez-Larios and Alaerts (2021) | Negative | Interoceptive (Breath) | Probe |
| Rout et al. (2024) | Negative | Interoceptive (Breath) | Self |
| van Son et al. (2019a) | Negative | Interoceptive (Breath) | Self |
| van Son et al. (2019b) | Negative | Interoceptive (Breath) | Self |

## Discussion

### Alpha oscillations and mind-wandering

The literature on mind-wandering faces a paradox: mind-wandering is associated with stronger alpha power in some studies but weaker alpha power in others. We reasoned that an inverted-U relationship between alpha power and mind-wandering could dissipate this paradox, with studies sampling different alpha power ranges (sides of the inverted-U) reporting opposite effects. We further hypothesized that whether participants’ eyes were open or closed could vary the range of alpha power explored by different studies. Accordingly, we found that mind-wandering was positively correlated with alpha power in the EO group, consistent with previous eyes-open studies using similar methods (Table 2, top) and the left side of the inverted-U. Eye closure (EC group) shifted alpha power into a higher range and reversed its correlation with mind-wandering, consistent with the right side of the inverted-U (Fig. 1).

Although previous eyes-closed studies reported a negative correlation (Table 2, bottom), whether it reflects the right side of the inverted-U or other methodological differences remains unclear. First, alpha power may show opposite effects in interoceptive and exteroceptive attention tasks (Cooper et al., 2003; Villena-González et al., 2017), though Shinagawa et al. (2025) found evidence against this. Second, eyes-closed studies assumed self-caught mind-wandering reports to be preceded by mind-wandering and followed by on-task states (Rout et al., 2024). However, if participants report mind-wandering only after becoming aware of it and returning on-task (Shinagawa et al., 2025), pre-report periods are mislabeled as “mind-wandering.” If participants immediately resume mind-wandering post-report, these periods are mislabeled as “on-task” (Rout et al., 2024). This mislabeling could reverse the sign of the observed correlation and limit interpretability of self-caught studies.

By using the same exteroceptive task and probe-caught experience-sampling in both eye states, our study demonstrates that the reversal of the correlation in the EC group arises from eye closure. Future research might test if this reversal replicates independent of eye closure by manipulating alpha power via perceptual deprivation (Ganzfeld: Miskovic et al., 2019), neurofeedback (Hanslmayr et al., 2005; Brickwedde et al., 2019), and neurostimulation (Vossen et al., 2015).

Taken together, our findings demonstrate that the paradoxical positive and negative correlations between alpha oscillations and mind-wandering reported previously, reflect two sides of an inverted-U relationship spanning alpha power ranging across eye states.

### Alpha oscillations and sleepiness

We first tested if subjective sleepiness was related to mind-wandering (Jubera-García et al., 2021). Participants mind-wandered more when they felt sleepier, and those with eyes closed showed both higher sleepiness and more frequent mind-wandering than those with eyes open. This is consistent with prior research showing that mind-wandering is more likely during states of low arousal reflected in subjective reports (sleepiness: Stawarczyk and D’Argembeau, 2016; Carciofo et al., 2014) and physiological metrics (reduced baseline pupil diameter: Unsworth and Robison, 2018; Stawarczyk et al., 2020; Andrillon et al., 2021; but see Smallwood et al., 2012; Groot et al., 2021; reduced stimulus-locked pupil dilation: Jubera-García et al., 2021; and strong EEG slow waves: Andrillon et al., 2021). Further, increasing sleepiness via sleep deprivation triggers mind-wandering (Poh et al., 2016; Walker and Trick, 2018; Yang et al., 2025) and, via pharmacology, increases attentional lapses (Pinggal et al., 2022), suggesting a causal link.

We next examined whether subjective sleepiness showed an inverted-U relationship with alpha power, as suggested by prior research. In the EO group, alpha power was positively correlated with sleepiness ratings, corresponding to the left side of the inverted-U. This aligns with previous eyes-open studies showing that alpha power increases with sleepiness measured directly through subjective ratings (Stampi et al., 1995; Cajochen et al., 1996; Kaida et al., 2006; Iemi et al., 2019), or indirectly through time-on-task (Benwell et al., 2019; Kopčanová et al., 2025). By contrast, in the EC group, alpha power was negatively correlated with sleepiness, consistent with the right side of the inverted-U. This replicates prior eyes-closed findings showing that alpha power decreases with increasing subjective sleepiness (Stampi et al., 1995; Strijkstra et al., 2003; Kaida et al., 2006; Snipes et al., 2024) and during sleep onset (“alpha dropout;” De Gennaro et al., 2001; Lacaux et al., 2024).

Taken together, these findings show that subjective reports of sleepiness and mind-wandering are behaviorally correlated and both exhibit an inverted-U relationship with alpha power.

### Alpha/Beta oscillations and reaction times

In our study, mean reaction times (RTs) didn’t differ between mind-wandering and on-task states in either group, inconsistent with our hypothesis (Leszczynski et al., 2017; Andrillon et al., 2021; but see Kam et al., 2011; Arnau et al., 2020). Note that our study did not distinguish between task-unrelated states with (i.e., mind-wandering) and without content (i.e., mind-blanking; Andrillon et al., 2024). These states may produce opposite effects on RTs (Andrillon et al., 2021; Munoz-Musat et al., 2025), so conflating them could yield null results.

We thus examined the relationship between prestimulus alpha power and RTs to replicate previous findings. In the EC group, we found that stronger alpha/beta power (in occipito-parietal electrodes) preceded faster (auditory) RTs. This is consistent with prior studies estimating oscillations from parietal and visual regions, and RTs in non-visual tasks (e.g. somatosensory: Linkenkaer-Hansen et al., 2004; ElShafei et al., 2022; auditory: Bollimunta et al., 2008; ElShafei et al., 2018; Brickwedde et al., 2025). However, the EO group showed mostly inconclusive evidence. This contrasts one study using source-level MEG, which likely boosts the signal-to-noise ratio of alpha oscillations, showing a similar relationship in both eye states (ElShafei et al., 2022). Thus, the effects on RTs, unlike subjective reports, are likely incompatible with an inverted-U across eye states. Importantly, because mean RTs didn’t track mind-wandering in our study, these results cannot provide evidence for or against the inverted-U model of mind-wandering.

### Mechanistic Interpretations

While this study doesn’t directly address why the inverted-U exists, by clarifying existing literature it provides a necessary first step toward understanding underlying mechanisms. We consider two mechanistic interpretations post-hoc.

First, alpha fluctuations during mind-wandering may reflect changes in arousal (see *Alpha oscillations and sleepiness*). Accordingly, the inverted-U we observed is compatible with mind-wandering occurring in states of decreased arousal indexed by (i) higher eyes-open alpha power (left side of the inverted-U) and (ii) lower eyes-closed alpha power (right side of the inverted-U). Decreased arousal is thought to induce a state of perceptual decoupling, reflected in dampened physiological responses to external sensory stimuli: for example, reduced event-related potentials at sleep onset (Harsh et al., 1994; Ogilvie, 2001; Andrillon and Kouider, 2020) and during mind-wandering (Baird et al., 2014), as well as reduced stimulus-locked pupil responses during mind-wandering (Jubera-García et al., 2020). Importantly, unlike selective attention, perceptual decoupling is thought to attenuate processing at all sensory sources, affecting task-related and task-unrelated information equally (Barron et al., 2011; Schooler et al., 2011). At the experiential level, this attenuation may yield mental states with either no content at all (mind-blanking) or with stimulus-independent content (i.e., unrelated to the immediate sensory environment; Boulakis et al., 2023).

Second, alpha fluctuations during mind-wandering may reflect shifts in selective attention between task-related and task-unrelated sensory information (Mazaheri et al., 2014; Morrow et al., 2023). According to functional inhibition, alpha oscillations modulate the excitability of neurons, locally enhancing or suppressing information processing within those neurons (Klimesch et al., 2007; Haegens et al., 2011; Iemi et al., 2022). Thus, the inverted-U may be compatible with (i) eyes-open alpha power inhibiting task-related sensory sources (likely reflecting mixed sources), shifting attention to non-auditory stimuli, and thereby increasing mind-wandering (left side of the inverted-U) and (ii) eyes-closed alpha power (likely reflecting a visual source) inhibiting task-unrelated sensory sources, shifting attention to auditory stimuli, and thereby decreasing mind-wandering (right side of the inverted-U).

An outstanding question is how to reconcile the inverted-U relationship between alpha power and arousal with the functional inhibition account. Strong alpha power in eyes-open states (left side of the inverted-U) reflects decreased arousal and plausibly induces perceptual decoupling through its known inhibitory effects (Iemi et al., 2019, 2022). However, the right side of the inverted-U challenges this view: in eyes-closed states, strong alpha power reflects increased arousal (Kaida et al., 2006) and excitability (de Graaf et al., 2017). Future work is needed to clarify the underlying neurophysiological mechanisms.

This study cannot adjudicate between these interpretations due to some limitations. First, the limited spatial resolution of our EEG recordings prevents us from determining whether eyes-open and eyes-closed alpha oscillations arise from different sensory sources (interpretation 2) or a global/common signal (interpretation 1). Second, because we did not probe content, we cannot determine whether changes in alpha power relate to mind-wandering about stimuli represented in the corresponding task-unrelated sensory sources (interpretation 2) or a mixture between mind-blanking and stimulus-independent mind-wandering (interpretation 1). Future studies could address this with higher-resolution recordings and more fine-grained experience-sampling methods.

In conclusion, these results demonstrate that the relationship between mind-wandering and alpha oscillations follows an inverted-U across eye states, with eye closure shifting alpha power into a higher range and reversing its correlation with mind-wandering. By considering eye state, this study clarifies contradictions in prior literature and specifies how ongoing neural oscillations shape subjective experience.

## Acknowledgements

We thank Barnard College students Emma Kornberg, Hiya Jain, Jena Mamdani, and Sophia Faisal for assistance with the EEG recording; Peter Balsam for providing resources, and Esperanza Jubera-García, Niko A. Busch, and Saskia Haegens for helpful feedback.

